# ModuleDiscoverer: Identification of regulatory modules in protein-protein interaction networks

**DOI:** 10.1101/119099

**Authors:** Sebastian Vlaic, Christian Tokarski-Schnelle, Mika Gustafsson, Uta Dahmen, Reinhard Guthke, Stefan Schuster

## Abstract

The identification of disease associated modules based on protein-protein interaction networks (PPINs) and gene expression data has provided new insights into the mechanistic nature of diverse diseases. A major problem hampering their identification is the detection of protein communities within large-scale, whole-genome PPINs. Current strategies solve the maximal clique enumeration (MCE) problem, i.e., the enumeration of all non-extendable groups of proteins, where each pair of proteins is connected by an edge. The MCE problem however is non-deterministic polynomial time hard and can thus be computationally overwhelming for large-scale, whole-genome PPINs.

We present ModuleDiscoverer, a novel approach for the identification of regulatory modules from PPINs in conjunction with gene-expression data. ModuleDiscoverer is a heuristic that approximates the community structure underlying PPINs. Based on a high-confidence PPIN of *Rattus norvegicus* and publicly available gene expression data we apply our algorithm to identify the regulatory module of a rat-model of diet induced non-alcoholic steatohepatitis (NASH). We validate the module using single-nucleotide polymorphism data from independent genome-wide association studies. Structural analysis of the module reveals 10 sub-modules. These sub-modules are associated with distinct biological functions and pathways that are relevant to the pathological and clinical situation in NASH.

ModuleDiscoverer is freely available upon request from the corresponding author.

## Introduction

Structural analysis of intracellular molecular networks has attracted ample interest over several decades [Albert, 2005]. This includes cellular networks such as protein interaction maps [Uetz *et al.*, 2000], metabolic networks [Ravasz *et al.*, 2002,Schuster *et al.*, 2002], transcriptional regulation maps [Lee *et al.*, 2002], signal transduction networks [Ma’ayan *et al.*, 2005] as well as functional association networks [Tong, 2004]. Recent advances in the field of network science have focused on the identification of modules within the organism specific interactome [Ivanov *et al.*, 2016]. The interactome captures interactions between all molecules of a cell [Sanchez *et al.*, 1999]. Common for biological networks, the interactome is represented as a graph composed of nodes denoting for cellular molecules that are connected by edges representing interactions between them. Within the interactome, modules are sub-graphs that can be linked to phenotypes. Up to date, the identification of modules has been applied mostly in the context of human diseases based on protein-protein interaction networks (PPINs) of *Homo sapiens*. Such disease modules have been successfully identified for, e.g., asthma [Sharma *et al.*, 2015], inflammatory and malignant diseases [Gustafsson *et al.*, 2014], obesity and type-2-diabetes (among others) [Barrenäs *et al.*, 2012]. They provide new in-depth insights into the underlying molecular mechanisms of diseases. For example, the asthma-associated module identified by [Sharma *et al.*, 2015] revealed three pathways that had not been directly associated with asthma before. Based on these findings a novel molecular mechanism in the regulation of inflammation in asthma was proposed.

**Figure 1.**
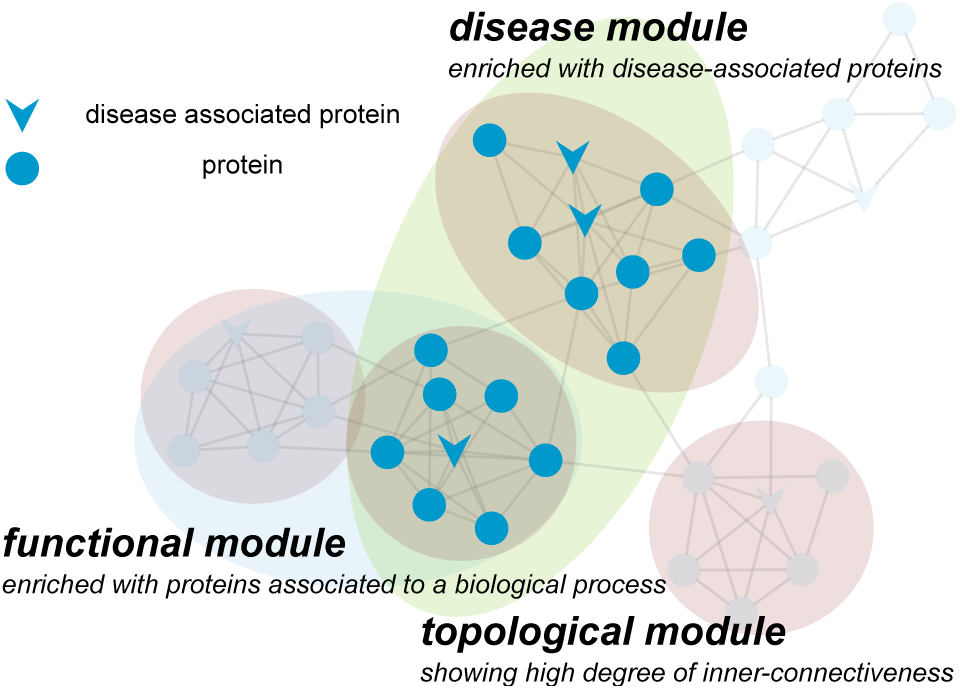
The concept of disease modules exemplified using a sample PPIN. One or more topological modules (highlighted red) contain proteins involved in similar biological processes forming functional modules (highlighted blue). A disease module (highlighted green) is a sub-network of proteins enriched with disease-relevant proteins, e.g., known disease associated proteins.

There are three fundamental assumptions to the identification of disease modules as summarized by [Barabási *et al.*, 2011] (Figure 1). Firstly, entities forming dense clusters within interactomes (topological modules) are involved in similar biological functions (functional modules). Secondly, molecules associated to the same disease, such as disease-associated proteins, tend to be located in close proximity within the network, which defines the disease module. Thirdly, disease modules and functional modules overlap. Thus, a disease can be seen as the breakdown of one or more connected functional modules.

A variety of approaches have been developed for the identification of disease modules. They can be roughly categorized into two different groups. On the one hand, there are algorithms that make use of known disease-associated molecules or genetic loci, the known interactome as well as some association function for the identification of disease modules and/or new disease-associated molecules [Oti *et al.*, 2006, George *et al.*, 2006,Köhler *et al.*, 2008,Ahn *et al.*, 2010,Ghiassian *et al.*, 2015]. For example, the disease module detection (DIAMOnD) algorithm [Ghiassian *et al.*, 2015] utilizes known disease-associated proteins (seed proteins) to identify proteins (DIAMOnD proteins) significantly connected to seed proteins. Iterative application of the algorithm results in a growing disease module with a ranked list of DIAMOnD proteins, i.e., candidate disease-associated proteins. On the other hand, there are algorithms that identify disease modules as well as disease-associated molecules “;ab initio ” based on the projection of omics data onto the interactome in conjunction with a community structure detecting algorithm [Barrenäs *et al.*, 2012, Gustafsson *et al.*, 2014, Zhang *et al.*, 2015]. Like topological modules, communities are groups of proteins with higher within-edge density compared to the edge density connecting them [Fortunato, 2010]. For example, the approach presented by [Barrenäs *et al.*, 2012] identifies protein communities by decomposition of the human PPIN into sub-graphs of maximal cliques. A clique is a sub-graph (a group of proteins) of the PPIN, where each pair of proteins is connected by an edge. A maximal clique is a clique that is not part of a larger clique. The regulatory module is then formed by the union of all maximal cliques that are significantly enriched with genes that are differentially expressed between samples of diseased and healthy subjects.

While these approaches have been presented specifically for the identification of disease-associated modules in *H. sapiens*, the idea is generalizable towards the detection of regulatory modules underlying an arbitrary phenotype of any organism. This can be of high interest, e.g., for the molecular characterization of animal models of human diseases, where the availability of human samples is limited. For example, the use of rodent models of fatty liver disease (FLD) provides a defined, rigorously controllable alternative to human liver tissue samples. They can be easily generated allowing for experiments with sufficient sample size to obtain statistically meaningful hypotheses. Furthermore, it is much easier to control environmental factors, which is of high importance for systematic investigation of multifactorial diseases like FLD. However, up to date there is no model that completely reflects all aspects of FLD [Imajo *et al.*, 2013]. Thus, identification of regulatory modules underlying animal models of FLD can provide important information regarding their relevance towards the human condition in FLD on the molecular level.

A major problem that hampers the efficient *ab initio* identification of regulatory modules is the detection of the community structure underlying PPINs. Algorithms solving the maximal clique enumeration (MCE) problem as utilized by [Barrenäs *et al.*, 2012] are *non-deterministic, polynomial time* (NP) -hard [Eblen *et al.*, 2012] and thus a computationally challenge regarding the processing of large-scale, genome-wide PPINs.

We present ModuleDiscoverer, a new approach to the *ab initio* identification of regulatory modules. ModuleDiscoverer is a heuristic that approximates the PPIN’s underlying community structure by iterative enumeration of cliques starting from random seed proteins in the network. To demonstrate its application we identify the regulatory module underlying a diet induced rat model of non-alcoholic steatohepatits (NASH), the severe form of the non-alcoholic fatty liver disease (NAFLD). The identified NASH- regulatory module is then validated using single nucleotide polymorphism (SNP) data of NAFLD from independent genome-wide association studies (GWASs) as well as known gene-to-disease relations. Moreover, we show that the identified regulatory module reflects histological and clinical parameters as reported by [Baumgardner *et al.*, 2008], who first introduced the animal model.

## 1 Methods

### Microarray data, pre-processing and differential gene expression analysis

Affymetrix microarray gene expression data of a rodent model of diet induced NASH published by [Baumgardner *et al.*, 2008] was downloaded from Gene Omnibus Express [Barrett *et al.*, 2013] (GSE8253). In brief, [Baumgardner *et al.*, 2008] obtained the animal model by overfeeding rodents with a high-fat diet based on 70% corn oil at moderate caloric excess (220*kcal* * *kg^−3/4^* * *day*^−1^ ∼ 17%) for 21 days via total enteral nutrition (TEN). They compared the treatment group against a control group of rats fed a diet based on 5% corn oil at normal caloric levels (187*kcal* * *kg*^−3/4^ * *day*^−1^) for 21 days via TEN. Gene expression in each experimental group was measured using three microarrays.

Affymetrix Rat Genome U34 arrays were annotated with custom chip definition files from Brainarray version 15 [Dai *et al.*, 2005]. Raw-data was pre-processed using RMA [Irizarry *et al.*, 2003]. Differential gene expression was assessed using *limma* [Ritchie *et al.*, 2015] with a p-value < 0.05 (additional file 2).

### SNP-gene-disease and gene-disease association data

Disease-to-SNP relations as well as curated disease-to-gene associations for *H. sapiens* were obtained from DisGeNET [Piñero *et al.*, 2015]. All text-mining based disease-to-SNP associations were removed. Furthermore, we removed all associations involving genes without an orthologue in *R. norvegicus*. Orthology information was obtained from the RGD [Shimoyama *et al.*, 2015]. For the disease-to-gene associations we created a “disease network” similar to [Goh *et al.*, 2007]. In this network, we connected two diseases (nodes) by an edge if they share >= 10 genes. Selecting the first neighbors of the terms “fatty liver” and “non-alcoholic fatty liver disease” yielded a list of 31 NAFLD-relevant diseases.

## 2 Results

### ModuleDiscoverer: detection of regulatory modules

The detection of regulatory modules is divided into 3 steps I - III (Figure 2). Starting with a PPIN (Figure 2, Input) the algorithm first approximates the underlying community structure by iterative enumeration of protein cliques from random seed proteins in the network (Figure 2, I). Next, DEGs obtained from high-throughput gene expression data in conjunction with sets of randomly sampled genes (Figure 2, Input) are used to calculate a p-value for each clique (Figure 2, II). Finally, significantly enriched cliques are assembled (Figure 2, III) resulting in the identified regulatory module (Figure 2, Output).

**Figure 2.**
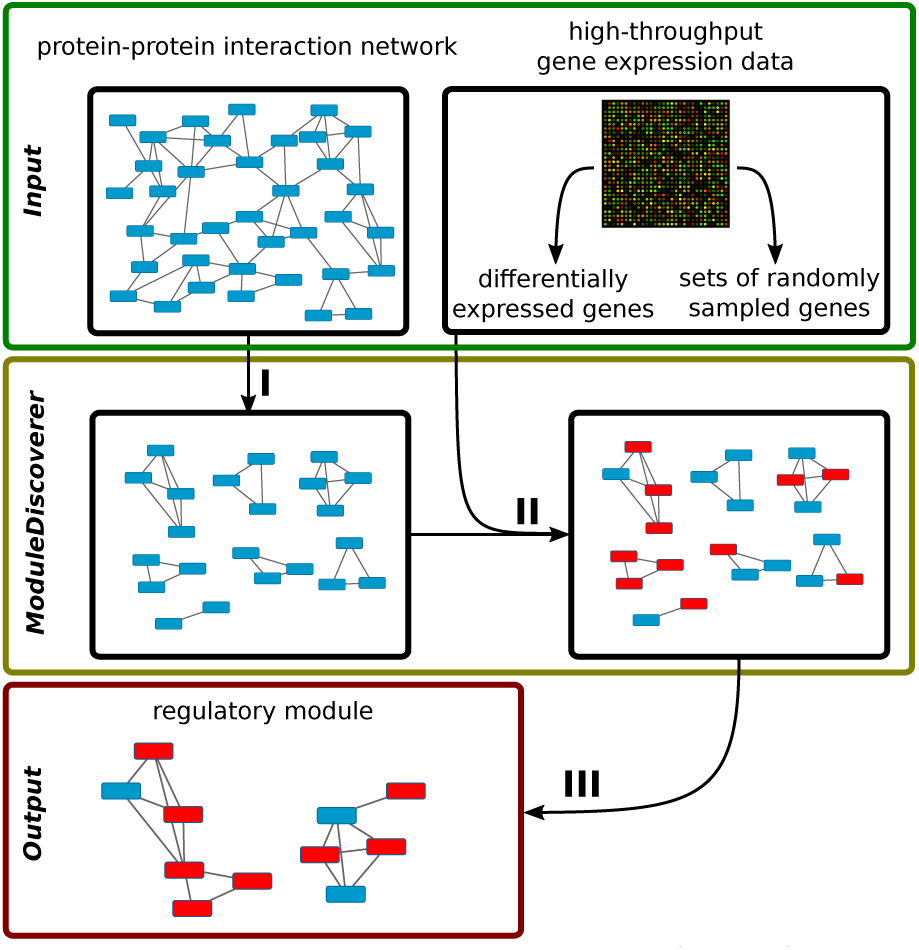
Given a PPIN and gene expression data (Input) the algorithm works in 3 steps. Step I) The community structure underlying the PPIN is approximated by the identification of protein cliques. Step II) Identification of cliques significantly enriched with DEGs. Step III) Assembly of the regulatory module based on the union of significantly enriched cliques.

### Step I: Approximation of the PPIN’s community structure

Approximation of the community structure underlying the PPIN (Figure 2, I) is composed of 3 phases: transformation, identification and extension. In brief, the PPIN is transformed into a graph with labeled nodes and edges (Figure 3, A - B). Starting from one or more random seed nodes the algorithm then identifies minimal cliques of size three (Figure 3, C - E). Finally, all minimal cliques are stepwise extended competing for the nodes in the network until no clique can be extended further (Figure 3, F).

**Figure 3.**
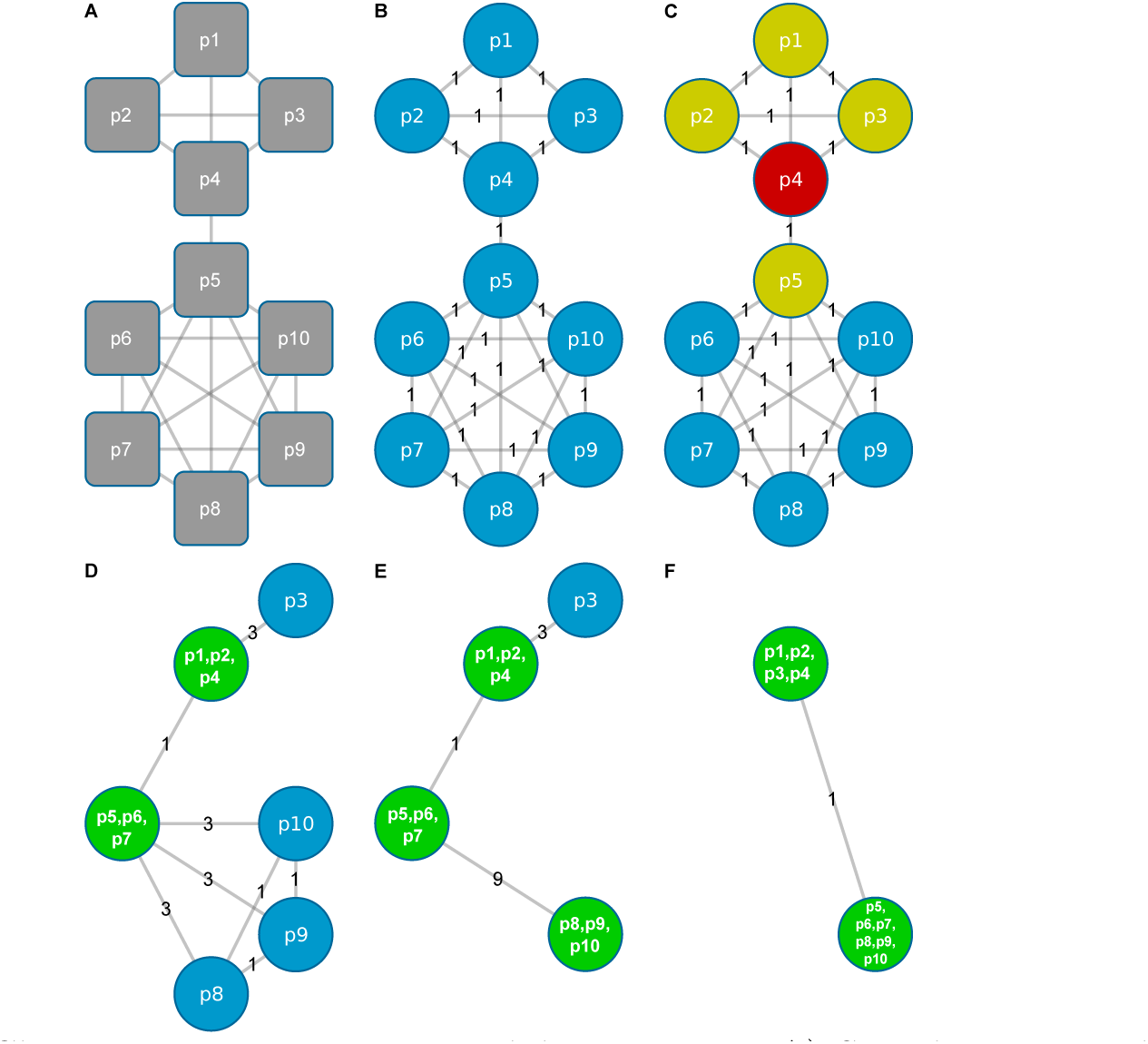
Clique enumeration using ModuleDiscoverer. A) Sample PPIN with 10 proteins and 26 known relations. B) Representation of the PPIN as a undirected labeled graph with each vertex representing one of the proteins in A). The edge weight denotes for the number of existing relations between its connecting nodes. C-F) Red vertices denote for seed nodes. Yellow vertices are first neighbors of seed nodes. Green vertices represent cliques. Their label represents clique forming proteins.

The number of seed nodes defines two strategies for the enumeration of cliques, the single-seed and the multi-seed approach. Notably, there are advantages as well as disadvantages for both strategies. Details on their performance as well as additional examples are provided in additional file 1. The results are summarized and evaluated in the discussion section. In the following example we will illustrate our approach showing one iteration of ModuleDiscoverer using three seed proteins (*p*4, *p*6 and *p*9).

### Phase 1 of Step I: Transformation of the PPIN into a labeled graph

Figure 3(A) shows a PPIN as provided by databases such as STRING [Szklarczyk *et al.*, 2015]. It consists of 10 nodes representing the proteins *p*1 to *p*10 and 26 connecting edges. These edges refer to prior-knowledge interactions between connected proteins. First, the network is transformed into an undirected labeled graph *G*(*V, E*) (Figure 3, B). The graph *G* consists of 10 vertices *V*(*G*) = {*v*_1_,…, *v*_10_} and 26 edges *E*(*G*) = {*e*_1_,…, *e_26_*}. Each vertex is labeled with one protein (*p*1 - *p*10). Notably, a vertex can be labeled with more than one protein. In such case, the proteins in the label form a clique in the PPIN (e.g., vertex *p*1,*p*2,*p*4 in Figure 3, D). Two vertices *v_x_* and *v_y_* (with *x*,*y* ∈ 1,…, 10 and *x* ≠ *y*) are connected by an edge if there is at least one known relation in the PPIN between the proteins represented by *v_x_* as well as the proteins represented by *v_y_*. The weight of the edge connecting *v_x_* and *v_y_* denotes for the number of relations between the proteins represented by *v_x_* and the proteins represented by *v_y_*. Initially, all edges have weight 1.

### Phase 2 of Step I: Identification of minimal cliques of size three

Starting with randomly selected seed proteins the algorithm first identifies minimal cliques of size three. A seed is dropped if it is not part of a minimal clique. In Figure 3 (C), we start with *p*4 (colored red) as a seed and search for any minimal clique of size three by exploring its neighbors (colored yellow) as well as their neighbors. The order in which vertices are explored is random. In our example, the first clique identified is formed by *p*1, *p2* and *p*4 and the corresponding vertices are merged into the vertex *p*1,*p*2,*p*4 (see Figure 3, D). Next, the weights of the edges are updated. In our example (Figure 3, D), the edge between *p*1, *p*2, *p*4 and *p*3 is now weighted 3, since the proteins *p*1,*p*2 and *p*4 are all connected to protein *p*3 (see Figure 2, A). The edge’s weight connecting *p*1,*p*2,*p*4 with *p*5 remains 1, since only *p*4 is connected to *p*5. Following the same strategy, the minimal clique *p*5,*p*6,*p*7 is identified starting from the seed *p*6 (Figure 3, D) while the seed *p*9 is merged with *p*8 and *p*10 into *p*8,*p*9,*p*10 (Figure 3, E). All edge weights are updated accordingly.

### Phase 3 of Step I: Extension of all minimal cliques

All minimal cliques of size three (Figure 3, E; green) are now iteratively extended in random order until they cannot be enlarged further. Once a node becomes part of a clique, it cannot become part of another clique, i.e., cliques compete for nodes in the graph. Starting from figure 3 (E), *p*1,*p*2,*p*4 is processed first. *p*1,*p*2,*p*4 is connected to *p*3 by an edge of weight 3. Thus, all proteins *p*1, *p*2 and *p*4 are connected to *p*3 (see Figure 3, A). Therefore, both vertices can be merged to form the new vertex *p*1,*p*2,*p*3,*p*4 (Figure 3, F). Next, the clique represented by *p5*,*p*6,*p*7 is processed. The edge connecting *p*5,*p*6,*p*7 with *p*8,*p*9,*p*10 has a weight of 9. This indicates that all proteins of *p*5, *p*6, *p*7 are connected with all proteins of *p*8, *p*9, *p*10. Therefore, both vertices are merged to form *p*5, *p*6, *p*7, *p*8, *p*9, *p*10 (Figure 3F). Finally, no clique can be enlarged any further. The algorithm terminates reporting two cliques, i.e., the clique formed by the proteins *p*1,…,*p*4 as well as the clique formed by the proteins *p*5,…,*p*10.

Phases 1 - 3 of step I of the algorithm are repeated for n iterations with random seed proteins in each iteration until the set of obtained cliques sufficiently approximates the community structure underlying the PPIN.

### Step II: Identification of significantly enriched cliques

In step II (Figure 2, II) all enumerated cliques are tested for their enrichment with phenotype-associated proteins, e.g., proteins corresponding to DEGs from high-throughput gene expression data (Figure 2, Input). The p-value for each clique is calculated using a permutation-based test [Ge *et al.*, 2003]. In detail, for a gene expression platform measuring *N* genes, with *D* ∈ *N* being the set of DEGs, the gene sets *B* are created, each containing |*D*| genes sampled from *N*. For each clique in *C*, the p-value *p_i,D_* of clique *c_i_* (i = 1,…, |*C*|) is calculated using the one-sided Fisher’s exact test. Accordingly, the p-value *p_i,b_* of clique *c_i_* is calculated for each gene set *b* in *B*. The final p-value 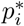 is then calculated according to equation 1.

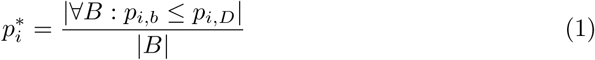

### Step III: Assembly of the regulatory module

Based on a user defined p-value cutoff we filter significantly enriched cliques. Since cliques can overlap in their proteins, the union of all significantly enriched cliques (Figure 2, III) results in a large regulatory module (Figure 2, Output). This module summarizes biological processes and molecular mechanisms underlying the respective phenotype.

### Reproducibility of regulatory modules

ModuleDiscoverer is a heuristic that approximates the underlying community structure. Since the exact solution is unknown, quality of the approximation cannot be assessed directly. Instead, we can test if additional iterations of the algorithm, i.e., the enumeration of more cliques, has a qualitative impact on the regulatory module in terms of additional nodes and edges. To this end, non-parametric bootstrapping sampling (with replacement) is applied to assess reproducibility of the regulatory module. Based on the results of n iterations of ModuleDiscoverer we create bootstrap samples of *n* iterations and identify the respective regulatory modules. Pairwise comparison of the regulatory modules in terms of shared edges and nodes then provides a distance between the two regulatory modules. The median of all distances divided by the average number of nodes and edges reflects the stability of the regulatory module. See additional file 1 section 1.4 for details.

## ModuleDiscoverer: application to biological data

To demonstrate the application of ModuleDiscoverer we used the PPIN of *R. norvegicus* in conjunction with gene expression data of a rat model of diet-induced NASH for the identification of a NASH-regulatory module. The results will be presented in three sections: (i) processing of the PPIN (Figure 2, I), (ii) identification of significantly enriched cliques based on high-throughput expression data (Figure 2, II) and, (iii) assembly of the regulatory module based on the union of all significantly enriched cliques (Figure 2, III). Finally, the NASH-regulatory module will be analyzed and validated.

### Processing of the PPIN

The PPIN of *R. norvegicus* (STRING, version 10) was filtered for high-confidence relations with a score > 0.7. This retained 15436 proteins connected by 474395 relations. Next, we used the single-seed approach of ModuleDiscoverer to enumerate maximal cliques using 2,000,000 iterations. This identified 1,494,126 maximal cliques in total, enclosing 185,178 unique maximal cliques. Additionally, we applied ModuleDiscoverer with 1,020,000 iterations using the multi-seed approach with 25 seed proteins per iteration. This resulted in 18,807,344 cliques in total enclosing 2,269,022 unique cliques.

### Identification of significantly enriched cliques

Based on the expression data we identified 286 DEGs (p-value < 0.05) out of 4590 EntrezGeneID-annotated genes on the microarray platform (see additional file 2). 10,000 data sets were created sampling 286 random genes out of 4590 genes in the statistical background. Finally, genes of all data sets were translated into EnsemblProteinIDs using the R-package *org.Rn.eg.db*.

P-value calculation according to equation 1 was performed for each clique satisfying the following two properties. First, at least one protein in the clique is associated to a DEG. Second, at least half of the proteins in the clique are associated to genes in the statistical background. For the p-value cutoff 0.01 we identified 696 significantly enriched cliques for the single-seed approach and 5386 significantly enriched cliques for the multi-seed approach. Notably, permutation-based calculated p-values were similar to p-values calculated using the one-sided Fisher's exact test (additional file 3).

### Assembly and analysis of the regulatory module

The assembled regulatory module (additional file 4) of the single-seed approach contains five sub-networks composed of 311 proteins connected by 3180 relations. 175 of the 311 proteins are associated to background genes and 60 are associated to DEGs. Similar, the regulatory module of the multi-seed approach contains five sub-networks composed of 415 proteins and 4975 relations in total (Figure 4). 210 of these 415 proteins are associated with background genes. 67 proteins are associated with DEGs. Both of the regulatory modules are significantly enriched (*p* < 10^−4^) with proteins associated to DEGs. Based on 100 bootstrap samples (see additional file 1 section 1.4) we found that both regulatory modules are reproducible with an average variability of less then 5% (additional file 5). Apart from a single edge the multi-seed regulatory module encloses the single-seed regulatory module. Thus, we decided to focus on the multi-seed regulatory module as an extension to the single-seed regulatory module.

**Figure 4.**
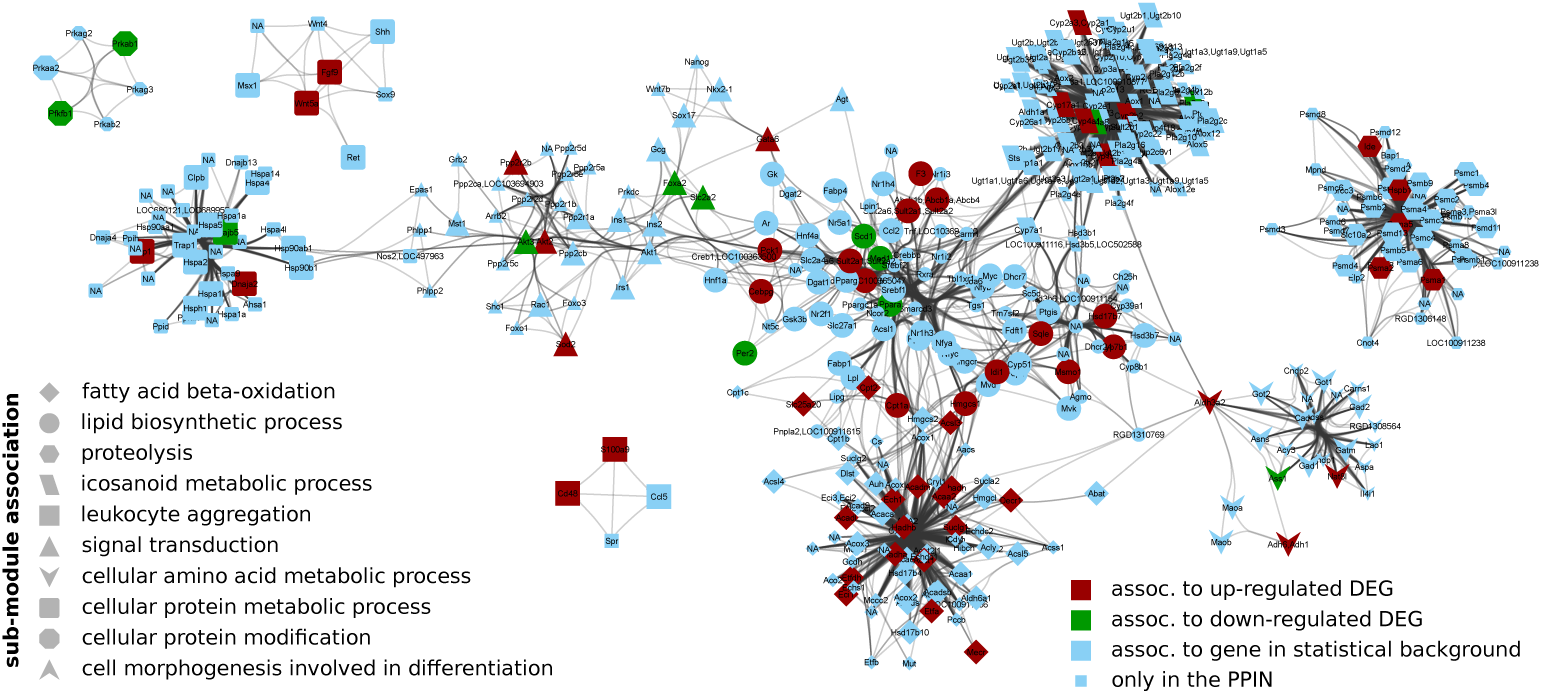
The identified NASH-regulatory module. Nodes (proteins) are labeled with the official gene symbol. Their membership in a sub-module is shape-coded.

Next, we identified pathways significantly enriched with proteins for the regulatory module shown in Figure 4. The results (additional file 6) highlighted NASH-relevant pathways such as fatty acid degradation and elongation, PPAR signaling pathway [Souza-Mello, 2015], arachidonic acid metabolism [Loomba *et al.*, 2015], the metabolism of diverse amino acids [Cheng *et al.*, 2012] as well as insulin signaling pathway [Nassir and Ibdah, 2014, Chitturi *et al.*, 2002]. Identification of sub-modules based on edge-betweenness [Newman and Girvan, 2003] in the network revealed 10 sub-modules. These sub-modules are sparsely connected with each other but densely connected within themselves. In Figure 4, the sub-module membership of each protein is shape-coded. We performed an enrichment analysis for the proteins of each sub-module to identify its potential biological function (additional file 7).

We found that the most central sub-module (Figure 4, circles) can be associated with lipid biosynthetic process. For example, the KEGG PPAR-signaling pathway is significantly enriched with proteins from the module. This pathway plays a key-role in the development of FLD by regulating the beta-oxidation of fatty acids, the activation of anti-inflammatory pathways and the interaction with insulin signaling [Pawlak *et al.*, 2015]. In agreement with these findings the sub-module is directly connected to sub-modules associated to fatty acid beta-oxidation (diamonds), icosanoid-metabolic processes (parallelogram) and cellular signal transduction such as the insulin signaling pathway (triangles). Another directly connected sub-module can be associated to the metabolism of cellular amino acids (V-shaped) such as alanine, aspartate and glutamate metabolism as well as phenylalanine, tyrosine and tryptophan metabolism.

The two remaining larger sub-modules can be associated to proteolysis (hexagons) and the metabolism of cellular proteins (round rectangle) with the latter being directly connected to the sub-module associated with signal transduction (triangles). The connection between cellular protein metabolic processes such as the response to unfolded proteins (see additional file 7, sub-module 8) and NAFLD as well as NASH has been studied extensively and is reviewed in [Henkel and Green, 2013].

### Literature validation of the regulatory module

Both NASH regulatory modules (single-seed and multi-seed) were validated using curated disease-to-SNP associations (see methods). Disease-to-SNP associations are based on DNA-sequence information. Thus, it can be considered independent from gene expression data, which was used to identify the regulatory module. We found that in contrast to the set of DEGs, both regulatory modules are significantly enriched (p-value < 0.05) with SNPs associated to NAFLD (additional file 8).

In addition to the validation based on SNP-information we used a list of curated disease-to-gene associations (see methods). The result of the enrichment analysis is summarized in Figure 5. Both regulatory modules show significantly enriched FLD-associated diseases such as obesity, (non-insulin dependent) diabetes mellitus type-2, liver carcinoma and insulin resistance. Notably, these disease-terms show a slight, but none significant enrichment for the set of DEGs. This demonstrates that the regulatory module is based on the integration of experimental data as well as PPI-knowledge.

**Figure 5.**
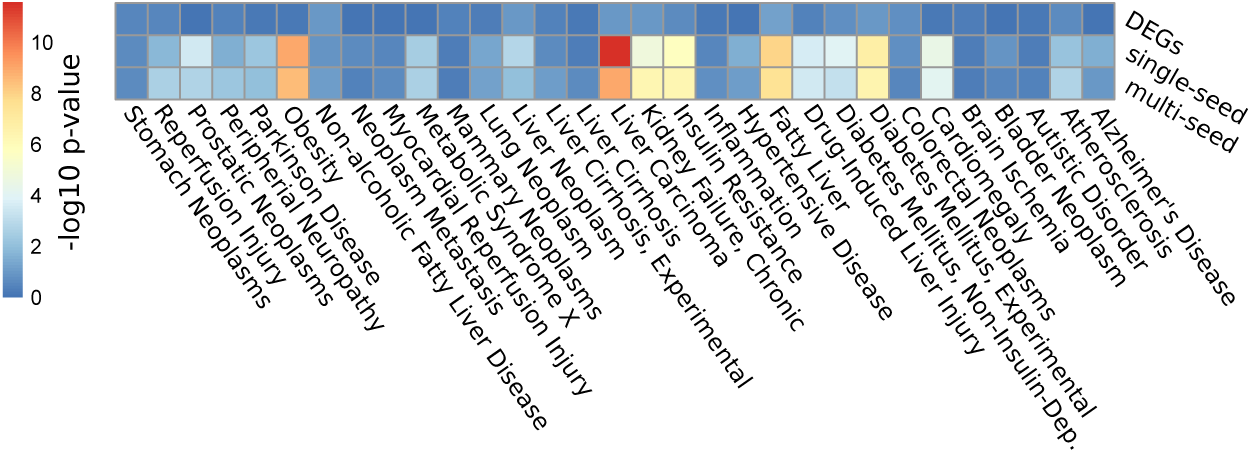
Enrichment of FLD-related diseases with proteins of the single-seed, the multi-seed regulatory module and proteins associated to DEGs. Higher values equal lower p-values.

## 3 Discussion

We have presented ModuleDiscoverer, an algorithm for the identification of regulatory modules based on large-scale, whole-genome PPINs and high-throughput gene expression data. To show applicability of the algorithm we identified a NASH-regulatory module for which we relied on the STRING resource only. STRING integrates information from a variety of resources, such as primary interaction databases, algorithms for interaction prediction, pathway databases, text-mining and knowledge transfer based on orthology. Reported relations are thus based on known physical interaction as well as associative information. To ensure quality of the relations we selected a high cutoff (>0.7) for the combined edge score. While this retains only high-confidence relations it may introduce knowledge bias. A yet to explore alternative might be the use of whole-genome gene regulatory networks (GRNs). Algorithms such as presented in [Altwasser *et al.*, 2012] are based on mathematical models that combine expression data and prior-knowledge interaction data. In such GRNs, relations denote for functional relationship between genes/proteins acting in common biological contexts, which equals networks derived from STRING [Szklarczyk *et al.*, 2015]. This corresponds to the idea of regulatory modules as shown in Figure 1.

To evaluate our algorithm (additional file 1) we used a small sub-network of the high-confidence PPIN of *R. norvegicus*. We showed that the single-seed approach as well as the multi-seed approach work well in principle and highlighted their advantages as well as disadvantages.

Using a single seed per iteration our algorithm will identify only maximal cliques. For large enough numbers of iterations the enumerated maximal cliques will cover all maximal cliques in the PPIN. These regulatory modules will be equal to the regulatory modules identified based on approaches utilizing MCE-problem solving algorithms such as *MACE* [Kazuhisa Makino, 2004], *c-isol* [Hüffner *et al.*, 2009], *CFinder* [Adamcsek *et al.*, 2006], *MCODE* [Bader and Hogue, 2003] and *igraph* [Csardi and Nepusz, 2006]. Importantly, these algorithms process PPINs systematically and thus efficiently enumerate all maximal cliques compared to the randomization-based approach of ModuleDiscoverer. However, given the PPIN we used in this study neither of these algorithms successfully terminated. A possible explanation for the large number of required iterations of ModuleDiscoverer could be the high overlap of maximal cliques in the PPIN. Within such networks, small maximal cliques that overlap larger maximal cliques are less frequently identified. Firstly, because a randomly selected seed protein is more likely a member of the larger clique. Secondly, during the iterative extension of cliques, the number of candidate proteins is higher. As a consequence, proteins appearing in only small cliques may likely be missed because the required number of iterations for their discovery is too high.

The multi-seed approach is intended to avoid the enumeration of mainly large maximal cliques in such PPINs. We found that multiple seeds per iteration decrease the probability for their enumeration and shift it towards overlapping smaller cliques. This effect can be explained by the competition of cliques for available nodes during their iterative extension. This leads to the breakdown of large cliques into smaller cliques reducing the probability for their enumeration. Consequently, proteins appearing in only small maximal cliques are more often identified. As a drawback, a high number of seeds leads to the excessive breakdown of large cliques. Thus, a large maximal clique, which is not significantly enriched with DEG-associated proteins may be selected because of all its small sub-cliques being significantly enriched. To this end, the final regulatory module has to be tested for its enrichment with DEG-associated proteins and eventually, the number of random seeds should be lowered.

Thus, in cases where large-scale, genome-wide PPINs cannot be processed by MCE-solving algorithms, i.e., the regulatory module based on the exact solution cannot be determined, the use of ModuleDiscoverer becomes favorable. In such situations it is advisable to look at the identified regulatory modules of both, the single-seed as well as the multi-seed approach. The single-seed based regulatory module is more consistent with results of MCE-based approaches. In turn, the multi-seed regulatory module will extend the single-seed based regulatory module with proteins that may have been missed due to a PPIN structure of highly overlapping maximal cliques.

Analysis of the identified NASH-regulatory module showed its association with FLD-related diseases. Notably, these diseases relate to the findings of [Baumgardner *et al.*, 2008].

The NASH-regulatory module (Figure 4) highlights the disease-term obesity as significantly enriched with proteins of the module (Figure 5). In agreement, [Baumgardner *et al.*, 2008] observed a significant increase in body weight in the treatment group compared to control (p≤0.05). Moreover, they reported a significant increase in fat mass as percentage of body weight between treatment and control reflecting adiposity. Additionally, serum leptin levels were observed to be significantly increased in the treatment group. The serum leptin level is a marker that positively correlates with obesity [Al Maskari and Alnaqdy, 2006].

Other significantly enriched disease terms include insulin resistance, diabetes mellitus type-2 and diabetes mellitus, experimental. [Baumgardner *et al.*, 2008] reported significantly increased serum insulin concentrations compared to control rats that were overfed with a high-fat 5% corn oil diet at (220*kcal* * kg^-3/4^ * *day^-1^* ~ 17%) for 21 days. They concluded that this observation points towards hyperinsulinemia, which can be due to insulin resistance and is often associated with type-2 diabetes.

Finally, we found the disease-term fatty liver significantly enriched in proteins of the module. [Baumgardner *et al.*, 2008] reported that histological examination of the liver samples showed steatosis, macrophage infiltration and focal necrosis in the treatment samples. This was accompanied by significantly elevated serum alanine aminotransferase (ALT) levels and significantly increased serum and liver triglyceride concentrations. Notably though, other inflammation associated scores such as hepatocellular ballooning and lobular inflammation/necrosis were reported to be elevated but not statistically significant. This could explain the non-significantly enriched disease-terms such as inflammation and liver-cirrhosis.

## 4 Conclusion

We presented ModuleDiscoverer, a heuristic approach for the identification of regulatory modules in large-scale, whole-genome PPINs. The application of ModuleDiscoverer becomes favorable with increasing size and density of PPINs. Compared to a MCE-based approach we demonstrated that ModuleDiscoverer identifies regulatory modules that can be identical (single-seed approach) or even more comprehensive (multi-seed approach). We successfully applied our algorithm to experimental data for the identification of the regulatory module underlying a rat model of diet induced NASH. The identified NASH-regulatory module is stable, biologically relevant and reflects experimental observations on the clinical and histological level. In contrast to the analysis based on DEGs alone, the NASH-regulatory module is enriched with proteins associated to NASH-relevant diseases such as fatty liver, obesity and insulin resistance. Furthermore, we found the regulatory module significantly enriched with NAFLD-associated SNPs derived from independent GWASs. Altogether, we consider ModuleDiscoverer a valuable tool in the identification of regulatory modules based on large-scale, whole-genome PPINs and high-throughput gene expression data.

## Acknowledgments

We thank Dr. Jens Schumacher (Institute of Stochastics) and Stefan Lang (Institute for Bioinformatics) from the Friedrich-Schiller-University Jena for helpful discussions.

## Additional Files

### Additional file 1 — Supplementary information

Supplementary information regarding the performance of ModuleDiscoverer.

### Additional file 2 — List of differentially expressed genes as reported by limma

For each probe set Id (ProbeSetId) the table shows the associated Entrez-Gene-Id (EntrezGeneId), the official gene symbol (GeneSymbol) and gene name (GeneName) as well as the log2 fold-change (logFC) and the computed p-value (P-Value).

### Additional file 3 — Comparing permutation-based p-values with p-values calculated using Fisher’s exact test

Fisher-exact test (green dot) and permutation based (blue line) p-value for each clique over its rank of the permutation based p-value. Left: plot for single-seed regulatory module; Right: plot for multi-seed regulatory module.

### Additional file 4 — Cytoscape session of single-seed and multiseed regulatory module

A cytoscape session (.cys) containing the identified single-seed as well as multi-seed regulatory module underlying the diet induced rat model of NASH. For each node, we also provide information regarding additional identifiers such as EntrezGeneId (internal), EnsemblProteinId (name), gene name (GENENAME) and symbol (SYMBOL) as well as information regarding the sub-module membership (BetweennessCommunityMembership), association to a differentially expressed gene (inForeground) and the actual log2 fold change (log2FC) or a background gene (inBackground) in the gene expression data of Baumgardener et al. [Baumgardner *et al.*, 2008].

### Additional file 5 — Stability of the identified regulatory module

Plot outlining the stability in terms of nodes (left) and edges (right) of the single-seed (top) and the multi-seed (down) regulatory module for increasing numbers of iterations. Both plots shows that with more iterations the stability of the identified regulatory modules increases. In other words, the use of additional iterations (and therefore the enumeration of additional cliques) has a negligible effect on the structure of the identified regulatory modules.

### Additional file 6 — KEGG Pathway and GO enrichment analysis results

Each sheet contains the list of significantly enriched (p-value < 0.05) KEGG pathway term ids (KEGGID) as well as GO-term ids (GO[BP,MF,CC]ID) for the three different ontologies (BP: biological process; MF: molecular function; CC: cellular compartment). For each ID, the term (Term), the p-value (Pvalue), the size (Size) as well as the number of proteins in the respective set of genes (Count) are provided.

### Additional file 7 — KEGG-pathway and GO-term enrichment results for the sub-modules of the multi-seed based regulatory module

Each sheet contains the list of significantly enriched (p-value < 0.05) KEGG pathway term ids (KEGGID) as well as GO-term ids (GO[BP,MF,CC]ID) for the three different ontologies (BP: biological process; MF: molecular function; CC: cellular compartment). For each term (Term), the p-value (Pvalue), the size (Size) as well as the number of proteins in the respective set of genes (Count) are provided.

### Additional file 8 — Results of the enrichment analysis using gene-disease as well as SNP-gene-disease associations from the DisGeNet database

For each disease id (DiseaseId), the tables outline the disease name (DiseaseName), the calculated p-value (P-value) as well as the number of genes (not) in the regulatory module ((n)M) and (not) associated to the disease ((n)D). Each table represents the enrichment results for the set of DEGs, all disease associated proteins in the single-seed or multi-seed regulatory module.

